# Trading Genome Vulnerability for Stable Genetic Inheritance: Active Retrotransposons Help Maintain Pericentromeric Heterochromatin Required for Faithful Cell Division

**DOI:** 10.1101/740753

**Authors:** Yajing Hao, Dongpeng Wang, Shuheng Wu, Xiao Li, Changwei Shao, Peng Zhang, Jia-Yu Chen, Do-Hwan Lim, Xiang-Dong Fu, Shunmin He, Runsheng Chen

**Affiliations:** Key Laboratory of RNA Biology, Institute of Biophysics, Chinese Academy of Sciences, 100101, China; University of Chinese Academy of Sciences, Beijing, 100101, China; Department of Cellular and Molecular Medicine, University of California, San Diego, La Jolla, CA 92093-0651, USA

## Abstract

Retrotransposons are extensively populated in vertebrate genomes, which, when active, are thought to cause genome instability with potential benefit to genome evolution. Retrotransposon-derived RNAs are also known to give rise to small endo-siRNAs to help maintain heterochromatin at their sites of transcription; however, as not all heterochromatic regions are equally active in transcription, it remains unclear how heterochromatin is maintained across the genome. Here, we attack these problems by defining the origins of repeat-derived RNAs and their specific chromatin registers in *Drosophila* S2 cells. We demonstrate that repeat RNAs are predominantly derived from active Gypsy elements, and upon their processing by Dicer-2, these endo-siRNAs act in *cis* and *trans* to help maintain pericentromeric heterochromatin. Remarkably, we show that synthetic repeat-derived siRNAs are sufficient to rescue Dicer-2 deficiency-induced defects in heterochromatin formation in interphase and chromosome segregation during mitosis, thus demonstrating that active retrotransposons are actually required for stable genetic inheritance.

## INTRODUCTION

Eukaryotic genomes contain both gene-rich and gene-poor regions, respectively corresponding to euchromatin and heterochromatin. Heterochromatin can be further divided into two classes: facultative, which is dynamic and marked by H3K27me3, and constitutive, which is largely stable and marked by H3K9me2/3(Grewal and Jia 2007). Constitutive heterochromatin is predominately associated with centromeric and pericentromeric regions, telomeres, and retrotransposons(Lippman et al. 2004). Constitutive heterochromatin plays important roles in genome organization in the nucleus(Avner and Heard 2001; Zhang et al. 2019), suppression of recombination to protect genome integrity(Grewal and Klar 1997), and stable genetic inheritance during development and differentiation(Allshire et al. 1995; Peters et al. 2001). These critical biological functions underscore the importance of repeat-rich sequences underneath constitutive heterochromatin, which used to be referred to as “junk” DNA in the genome. In fact, besides their potential contribution to genome evolution, it has been unclear whether active retrotransposons have any immediate benefit to an organism.

Regarding the formation and maintenance of constitutive heterochromatin (thereafter generally referred to as heterochromatin), our current knowledge is largely derived from elegant genetic and biochemical studies in fission yeast and *Drosophila melanogaster*(Tschiersch et al. 1994; Grewal and Jia 2007; Holoch and Moazed 2015b). The most striking aspect of the emerging theme is that transcription is required for initiating heterochromatin formation, even though the eventual fate is to shut down transcription. In fission yeast, initial repeat-derived transcripts are amplified by an RNA-dependent RNA polymerase (RdRP). Resultant double-stranded RNAs are next processed by Dicer to produce small interfering RNAs (siRNAs), which are then loaded onto Ago1 to form the RNA-induced transcription silencing (RITS) complex to target nascent repeat RNA. RITS recruits a key histone methyltransferase Clr4 (Su(var)3-9 in flies and SUV39H1 in humans) to generate H3K9me2/3, which then attracts its reader Swi6 (HP1 in flies and humans), together inducing a series of RNA-protein and protein-protein interactions to mediate both initial deposition and spreading of H3K9me2/3 to neighboring sequences(Volpe et al. 2002; Verdel et al. 2004). *Drosophila melanogaster* appears to follow a similar scheme except using the piRNA system to process and amplify repeat-derived RNAs to establish heterochromatin to actively repress retrotransposition in the germline(Vagin et al. 2006; Halic and Moazed 2009; Muerdter et al. 2013; Iwasaki et al. 2015).

While the general conceptual framework for heterochromatin formation has been well established, there are multiple puzzles that remain to be solved. First, heterochromatin is still dynamic, rather than completely inert, raising the question of how transcription is restarted and whether heterochromatin maintenance depends on local transcription in all regions in need to be patched up. Second, in principle, some repeat-derived RNAs may also be capable of acting in *trans*, as suggested by a recent analysis of crosstalk between a reporter gene with one copy localized near a pericentromeric region of one chromosome and the other in its native euchromatic context of another chromosome in fission yeast(Yu et al. 2018). However, it remains unclear whether this principle generally applies to heterochromatin maintenance on all pericentromeric regions, and if so, what is the relative contribution of *cis*-versus *trans-*acting RNAs to such maintenance. Third, both fission and *Drosophila* germ cells are equipped with an RNA amplification system (RNA-dependent RNA polymerase in fission yeast and the ping-pong cycle in the fly), but such system is lacking in somatic cells of flies and mammals(Stein et al. 2003). The question is how somatic cells meet this supply/demand dilemma. Forth, fission yeast uses siRNA to drive heterochromatin formation and maintenance, but fly enlists the piRNA pathway for such purpose in the germline. This raises the question of whether different organisms may use distinct machineries for processing repeat-derived RNAs. Last, but not least, Dicer deficiency has been reported to cause chromosome mis-segregation during mitosis in both flies and mammals(Pek and Kai 2011; Huang et al. 2015), but it has remained unclear whether such dramatic phenotype results from impaired production of some sort of endo-siRNAs or other function(s) of Dicer in the nucleus.

Addressing the above questions would require critical information on where repeat-rich transcripts are generated and where these repeat RNAs target specific loci in the genome, which has been a challenging problem. Recent technological innovations have made it possible to comprehensively elucidate the RNA-chromatin interactome, which has been applied to multiple cell types, including *Drosophila* S2 cells(Li et al. 2017; Sridhar et al. 2017; Bell et al. 2018), but rich information on repeat-derived RNAs has not yet been explored. We have now utilized this information to define the interaction of repeat-derived RNAs with chromatin across the fly genome. Our analysis reveals that endo-siRNAs are mostly derived from the Gypsy family of retrotransposons, which are able to act in *cis* and *trans* to balance local transcription to help maintain pericentromeric heterochromatin. Strikingly, we show that a pool of synthetic endo-siRNA mimics is sufficient to rescue chromosome segregation defects in Dicer-deficient cells. These findings reveal that active retrotransposons are functionally required for maintaining heterochromatin to ensure stable genetic inheritance during cell cycle.

## RESULTS

### Strategy for genome-wide assignment of multi-mapped RNA and DNA reads

We recently developed a technology called global RNA-DNA interaction sequencing (GRID-seq) to detect chromatin-associated RNAs and their respective binding sites genome-wide. GRID-seq employs a bivalent linker to ligate to RNA in one end and fragmented DNA in the other end on fixed nuclei followed by selection and cleavage of linker ligated products with a type IIS restriction enzyme (Mme I) to generate “mated” RNA and DNA (both ∼20nt in length) for deep sequencing(Li et al. 2017). To control for the specificity in RNA-DNA mating, we also generated a GRID-seq library on mixed human and fly cells, thus enabling the construction of a critical background model by using cross-species reads (i.e. fly RNA ligated to human DNA and vice versa). Using these datasets, we previously explored uniquely mapped RNA-DNA mates to reveal nascent RNA-covered “transcription hubs” where specific promoters and enhancers are interconnected in the nucleus(Li et al. 2017). Given such high-quality data, we herein explored the biological significance of multi-mapped RNA and DNA reads in *Drosophila* S2 cells, taking advantage of high density reads on the much smaller genome of fly relative to humans.

The two independently generated GRID-seq libraries on S2 cells (Supplementary Table 1) were of high global concordance between RNA or DNA reads (Supplementary Fig. 1A). Interestingly, both replicates showed a high percentage (65% to 75%) of RNA reads that were mapped to repeat-derived transcripts, but a much smaller fraction (10% to 20%) of DNA reads were assigned to repeat-rich genomic loci (Supplementary Fig. 1B). To fully utilize these multi-mapped RNA and DNA reads, we took a previously established ShortStack strategy to make assignment to specific transcripts and genomic locations based on the local density of uniquely mapped reads(Axtell 2013). As illustrated (Fig. 1A), we first aligned each RNA read to genome and assigned it to annotated unique RNA transcripts according to the FlyBase database(Drysdale 2008) or repeat-derived transcripts based on the RepeatMasker database (http://www.repeatmasker.org), and each DNA read to genomic fragments(Roberts et al. 2015), generated by Alu I, a restriction enzyme used to fragment the fly genome during library construction. For multi-mapped reads, we distributed them to individual RNA transcripts or DNA fragments according to the relative density of uniquely mapped reads (Supplementary Fig. 1C).

**Figure 1.**
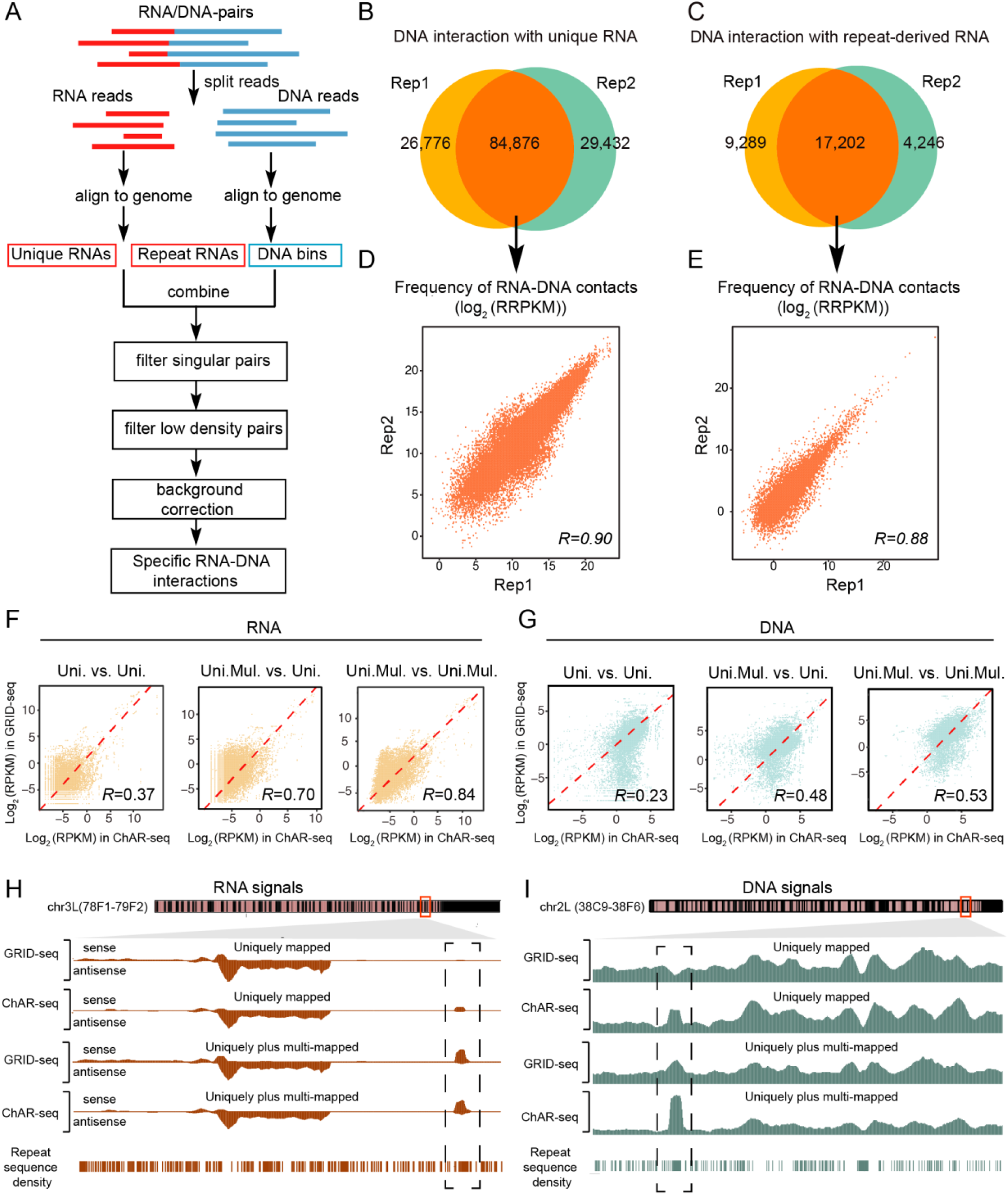
Strategy for assigning multi-mapped RNA and DNA reads. (A) Schematic presentation of the GRID-seq data processing pipeline. (B and C) Overlapped RNA-DNA mates containing annotated unique (B) or repeat-derived (C) RNA between the two independent GRID-seq libraries.(D and E) Quantitative analysis of commonly identified RNA-DNA mates containing annotated unique RNA (D) or repeat-derived RNA (E) between the two independent GRID-seq libraries. RRPKM: Read counts per kilobase of RNA per kilobase of DNA per million. (F and G) Comparison between GRID-seq and ChAR-seq with uniquely or uniquely plus multi-mapped RNA (F) or DNA (G) in repeat-enriched Alu I-generated DNA bins. Uni.: uniquely mapped reads. Uni.Mul.: uniquely plus multi-mapped reads. (H and I) Transcribed RNA signals (H) or RNA-contacted DNA signals (I) obtained with different mapping strategies on representative genomic regions.

We next filtered out singular RNA-DNA mates and low-density mates below a set threshold according to the Poisson distribution of all mates across the genome (see Online METHODS). Additionally, we also subtracted the background based on human RNA signals mapped to fly DNA loci from our fly/human GRID-seq library. The density of such background reads is quite significant in many accessible chromatin regions (Supplementary Fig. 1D), as detailed earlier(Li et al. 2017). After these data processing steps, retained RNA-DNA mates show high consistency between the two independent GRID-seq libraries, as indicated by predominant common mates associated with both annotated unique RNAs (Fig. 1B) and repeat-derived transcripts (Fig. 1C). This consistency is also reflected at the quantitative levels of individual common RNA-DNA mates (Fig. 1D,1E), thus enabling us to rely on these common mates to generate the final RNA-DNA interactome for downstream analysis. Notably, after assigning multi-mapped RNA and DNA reads, most gaps around repeat-rich DNA regions were “filled” to the similar levels, as compared to adjacent unique regions across the fly genome (Supplementary Fig. 2A).

### Validation of mapping results with an independent dataset

It is critical to validate our mapping strategy, even though ShortStack has been generally accepted as a strategy to dynamically assign multi-mapped reads in a given genome. For this purpose, we utilized the data generated by ChAR-seq, a strategy similar to GRID-seq except longer RNA and DNA reads were generated by sonication after linker ligation (Supplementary Fig. 2B)(Bell et al. 2018). Compared to GRID-seq that generates predominantly mated RNA-DNA pairs, ChAR-seq tends to trade off relative longer reads with a large fraction of unmated RNA or DNA reads from sequenced libraries. Moreover, it was not optimal to use the ChAR-seq data in the first place for several reasons: (i) GRID-seq libraries were generated on *Drosophila* S2 cells where many other types of genomic data are available for comparison (see Supplementary Table 2), whereas ChAR-seq libraries were produced on a less commonly used *Drosophila* cell line (CME-W1-cl8+), (ii) the vast majority of ChAR-seq reads was from a single library (see Supplementary Table 1), thus prohibitive to assessing internal data reproducibility, and most importantly, (iii) one of our GRID-seq libraries was constructed on mixed fly and human cells, thus permitting the use of cross-species RNA-DNA mates to build a background for non-specific RNA-DNA interactions, which is missing from the existing ChAR-seq libraries. Nevertheless, the available ChAR-seq data with longer RNA and DNA reads provided an independent dataset to evaluate our strategy for assigning multi-mapped RNA and DNA reads, despite the fact that GRID-seq and ChAR-seq libraries were derived from different fly cell types.

We first compared between ChAR-seq and GRID-seq data, observing an overall high Spearman correlation (R=0.75 for RNA reads; R=0.62 for DNA reads) across the reference fly genome (Supplementary Fig. 2C). However, when focused on repeat-enriched Alu I-generated DNA bins, the correlation was quite modest at the levels of both RNA (left panel in Fig. 1F, R=0.37) and DNA (left panel in Fig. 1G, R=0.23). Notably, a population of DNA reads (distributed in lower right in Fig. 1G) was scored by ChAR-seq, but less by GRID-seq, likely due to the higher mapping power of the former. Interestingly, the correlation was dramatically improved when comparing uniquely mapped RNA or DNA reads from ChAR-seq with uniquely plus multi-mapped RNA or DNA reads from GRID-seq (middle panels in Fig. 1F, R=0.70 for RNA and Fig. 1G, R=0.48 for DNA), suggesting that many assigned multi-mapped GRID-seq reads matched uniquely mapped ChAR-seq reads. As expected, after also assigning multi-mapped reads from ChAR-seq using ShortStack, the correlation was further improved when compared between uniquely plus multi-mapped RNA or DNA reads in both cases (right panels in Fig. 1F, R=0.84 for RNA and Fig. 1G, R=0.53 for DNA), which becomes comparable to the overall correlation across the genome (see Supplementary Fig. 2C, the remaining differences likely, at least in part, result from different cell types used to construct GRID-seq and ChAR-seq libraries). The progressive improvement is also evident on representative genomic regions, showing gained RNA (Fig. 1H) or DNA (Fig. 1I) signals, each in a repeat-rich region (dashed boxes), both of which were detectable with uniquely mapped ChAR-seq reads, missing from uniquely mapped GRID-seq reads, and then became visible after assigning multi-mapped GRID-seq reads. Together, these data validate our computational strategy to assign multi-mapped RNA and DNA reads to the genome.

### Preferential interaction of distinct RNA classes with eu-versus hetero-chromatin

Having maximally utilized both uniquely and multi-mapped RNA and DNA reads from our GRID-seq libraries, we next wished to investigate how different classes of RNA might differentially interact with DNA in the fly genome. We were first focused on annotated non-repeat RNAs with respect to their interactions with the genome. As reported earlier(Li et al. 2017), among 4,856 non-repeat RNAs associated with chromatin, most showed interactions with DNA near their sites of transcription (Supplementary Fig. 3A). Upon further subdividing different classes of such RNAs, we noted that most annotated snoRNAs and snRNAs were not only expressed but also extensively engaged in interactions with chromatin in *Drosophila* S2 cells, whereas virtually no pre-miRNA was associated with chromatin (Supplementary Fig. 3B). The intermediate levels of lncRNAs, mRNAs, and tRNAs on chromatin probably reflected wide expression ranges of these RNA species, and thus, only those with sufficient expression were detectable on chromatin.

In contrast to the chromatin binding patterns of annotated non-repeat RNAs, we also identified 230 repeat-derived RNA species that showed significant interactions with chromatin, which include all rRNAs, satellite DNA-transcribed RNAs (including those from simple repeats), and >70% of LINE and LTR-derived RNA species (Supplementary Fig. 3C). Percentage wise, the majority of RNA species was from LTR (45%), simple repeat (28%), and LINE (14%) classes (Fig. 2A). We then displayed these 230 repeat-derived RNAs across the fly genome and observed two general patterns, one scattering across the genome and the other concentrating on pericentromeric regions in chromosome 2 and 3 (large dashed boxes in Fig. 2B). A subset of these repeat-derived RNAs also bound chromosome 4 (small dashed boxes in Fig. 2B), which is known to be predominantly heterochromatic(Sun et al. 2000).

**Figure 2.**
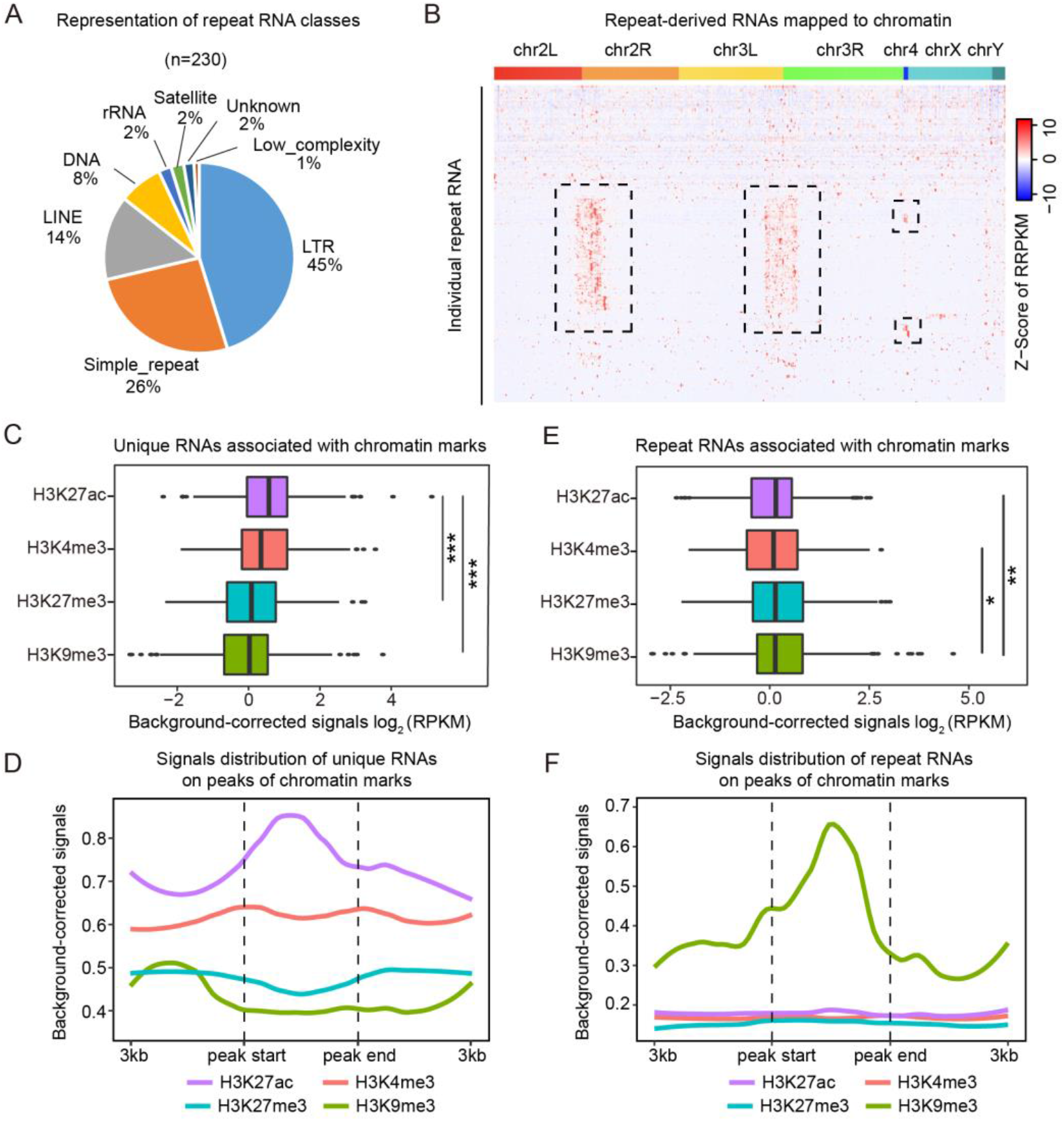
Interaction of distinct RNA classes with eu-versus hetero-chromatin. (A) Different repeat RNA classes represented by 230 DNA bound repeat-derived RNAs. (B) Heatmap showing the distribution of individual 230 repeat-derived RNAs across the *Drosophila* genome in S2 cells. Row: Individual chromatin-associated repeat-derived RNAs. Column: AluI DNA bins. Boxed regions: Repeat-derived RNAs that showed preferential binding to constitutive heterochromatin in pericentromeric regions. (C) Association of chromatin marks with background-corrected signals of unique RNAs. *p<0.05, **p<0.01, *** p<0.001 (unpaired Student’s t-test). (D) Distribution of unique RNA interaction signals around chromatin mark peaks. (E) Association of chromatin marks with background-corrected signals of repeat-derived RNAs. *p<0.05, **p<0.01 (unpaired Student’s t-test). (F) Distribution of repeat RNA interaction signals around chromatin mark peaks.

We next linked RNA-DNA interactions to critical chromatin features based on various published epigenetic profiles (Supplementary Table 2). As expected, annotated non-repeat RNAs showed the highest association with H3K27ac (Fig. 2C), exhibiting coincidental peak summits (Fig. 2D). In contrast, repeat-derived RNAs displayed the greatest preference for H3K9me3 (Fig. 2E,F). Conversely, we sorted the restriction enzyme Alu I-generated DNA fragments according to the levels of associated repeat-derived RNAs, observing that, among top 1,000 repeat RNA-associated DNA bins, H3K9me3 was the dominant signal on these DNA regions (Supplementary Fig. 3D). Together, these data suggest a general trend in which annotated non-repeat RNAs tend to interact with euchromatic regions marked by H3K27ac, whereas repeat-derived RNAs prefer to associate with heterochromatin marked by H3K9me3. While this is somewhat anticipated from the vast literature, we now obtained critical information on which specific repeat-derived RNAs are more prevalent than others in interacting with specific heterochromatic regions in the fly genome. This established the critical foundation to investigate their relative contributions to the initiation and/or maintenance of heterochromatin in *Drosophila* cells.

### Prevalent association of Gypsy-derived RNAs with constitutive heterochromatin

We next asked which repeat-derived RNAs were more prevalent than others on heterochromatin. We intersected the density of individual repeat-derived RNAs with markers for constitutive heterochromatin characterized by the coordinated ChIP-seq signals for H3K9me3 and its reader HP1 (Fig. 3A). By determining the co-localization coefficient for each of the 230 repeat-derived RNAs between their DNA interaction frequencies and relative densities of H3K9me3 and HP1 signals in 1Mb DNA bins of the fly genome, we identified 79 repeat-derived RNAs, including two rRNAs, that showed the *Pearson* correlation coefficient of >0.3 (red dots in Fig. 3A). The association of rRNAs with heterochromatin agrees with a recent observation that transcriptionally inert centromere-proximal regions tend to be organized around the nucleolus in 3D genome(Quinodoz et al. 2017). However, since rRNAs are assembled into ribosomes, rather than processed into small RNAs, it is unlikely that they contribute to heterochromatin functions. Excluding rRNAs, we named the rest of heterochromatin-enriched RNAs as CHARRs (<u>C</u>onstitutive <u>H</u>eterochromatin-<u>A</u>ssociated <u>R</u>epeats-derived <u>R</u>NA<u>s</u>). These CHARRs appear to show exclusive association with constitutive heterochromatin, as none of them exhibited significant co-localization with the facultative chromatin marker H3K27me3 (Supplementary Fig. 4A), as illustrated with three specific CHARRs (Supplementary Fig. 4B). Furthermore, these CHARRs showed little association with MSL (Supplementary Fig. 4C,D), a key component of the silencing complex involved in X inactivation in *Drosophila*, consistent with the nature of predominant facultative heterochromatin formed on X-inactivation (Xi)(Baker et al. 1994; Franke and Baker 1999).

**Figure 3.**
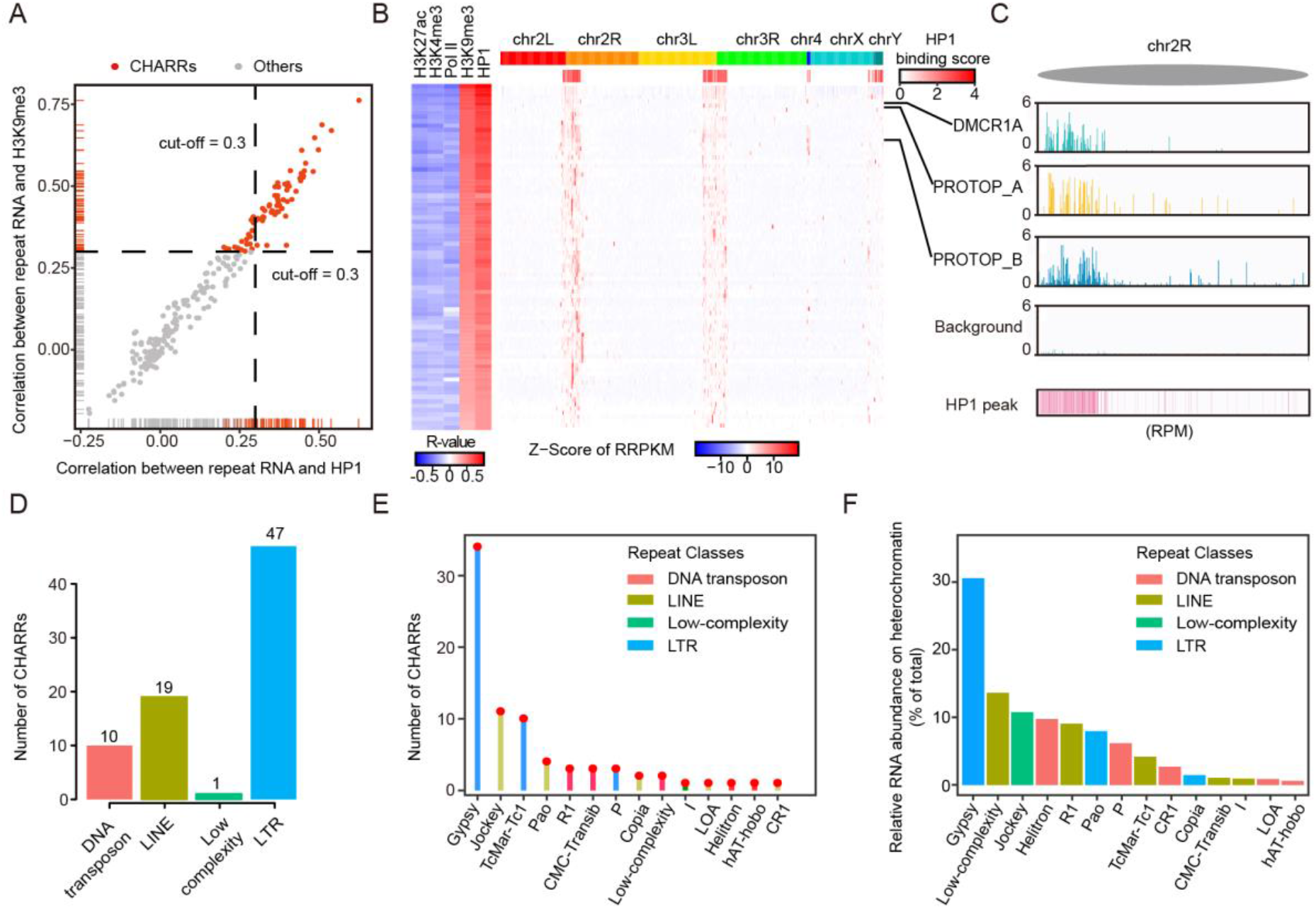
Prevalent association of Gypsy-derived RNAs with constitutive heterochromatin. (A) Scatterplot of co-localization coefficients between repeat-derived RNA signals on DNA and the levels of H3K9me3 (y-axis) or HP1 (x-axis) in S2 cells. A threshold of 0.3 was chosen for both chromatin marks (red lines). Dashed lines were used to differentiate CHARRs (red dots) from other repeat-derived RNAs (grey dots). (B) Left: Heatmap of co-localization of each CHARR with individual chromatin marks or Pol II ChIP-seq signals. Right: Heatmap of chromatin interaction of each CHARR with Alu I-generated DNA bins, which were merged from 100 continuous Alu I DNA bins. HP1 binding signals in these DNA bins are displayed below the schematic presentation of Drosophila chromosomes. Three representative CHARRs are labeled on the right. (C) Chromatin interaction signals of three representative CHARRs, DMCR1A, PROTOP_A and PROTOP_B on a genomic region in chromosome 2R in comparison with non-specific background based on human RNA mapped to fly DNA and HP1 binding density. All signals were scaled to reads per million. (D and E) The number of repeat classes (D) or sub-families (E) associated with CHARRs. (F) The relative RNA abundance (% of total) of CHARRs on constitutive heterochromatin and specific subfamilies they belong to. Colors show the RNA classes individual CHARR subfamilies belong to.

The 77 CHARRs we identified correspond to those predominantly associated with pericentromere of chromosome 2 and 3 where their interactions with DNA positively correlated to heterochromatin markers (e.g. H3K9me3 and HP1) and negatively to euchromatin markers (e.g. H3K27ac and H3K4me3) as well as RNAP II ChIP-seq signals (Fig. 3B). This is further highlighted with 3 specific repeat-derived RNAs (e.g. DMCR1A,PRTOP_A and PRTOP_B) on the right arm of chromosome 2 (chr2R, Fig. 3C). We next determined RNA class, genomic origins, and relative abundance for each CHARR. Interestingly, the largest RNA class of identified CHARRs corresponds to LTR (n=47), while the second largest class to LINE repeats (n=19) (Fig. 3D). The majority of RNA species from these two classes of retrotransposons belongs to the Gypsy and Jockey subfamilies, respectively (Fig. 3E). Most importantly, by ranking individual CHARRs according to their relative abundance on heterochromatin and summing the collective abundance according to specific RNA subclasses, we found that active Gypsy family members were top contributors to the overall RNA signals on heterochromatin (Fig. 3F). This suggests a major role of Gypsy-derived RNAs in heterochromatin formation/maintenance in *Drosophila* S2 cells.

### *Cis*- and *trans*-acting repeat-derived RNAs on chromatin

The RNA-DNA interactome permitted for the first time to determine both the source of repeat-derived RNAs and their registers on specific chromatin regions. Given the predominant mode of annotated non-repeat RNAs that act in *cis* (*cis* defined by mated RNA and DNA reads mapped to the same chromosomes, as supposed to trans defined by mated RNA and DNA reads mapped to different chromosomes), we asked whether this also applied to repeat-derived RNAs. We noted that rRNA-derived RNAs all interacted with heterochromatin regions near the loci of their transcription where CHARRs also predominantly bound (Supplementary Fig. 5A,B). This suggests multiple active rRNA transcription sites are in close spatial proximity with adjacent pericentromeric regions, as recently observed based on proximity ligation(Quinodoz et al. 2017). Importantly, we also identified CHARRs on multiple non-pericentromeric regions, suggesting their potential interactions with chromatin in both *cis*- and *trans*-modes.

We therefore determined the origins of CHARRs and their collective interactions with chromatin at the chromosomal levels. This analysis revealed their extensive chromatin interactions not only within the same chromosomes, but also across different chromosomes (Fig. 4A), suggesting a significant degree of *trans*-interactions. This prompted us to examine individual CHARRs to segregate their *cis*- and *trans*-interactions by first assigning their origins of transcription, and then determining their linkage to DNA on the same (intra) or different (inter) chromosomes. Interestingly, we found that about half (46.8%) of the CHARRs were predominantly engaged in intra-chromosomal interactions, whereas the other half (53.2%) were actively involved in both intra- and inter-chromosomal interactions (Fig. 4B). This is further illustrated with 3 representative CHARRs. Specifically, Gypsy12_LTR preferentially interacted with DNA on the same chromosome (Fig. 4C); Gypsy4_I-int was engaged in both intra- and inter-chromosomal interactions (Fig. 4D); and FW_DM seemed to mainly act in *trans* on other chromosomes (Fig. 4E). It is also interesting to note that individual CHARRs all selectively bound Hi-C defined “B” domains in pericentromeric regions (Supplementary Fig. 5C,D).

**Figure 4.**
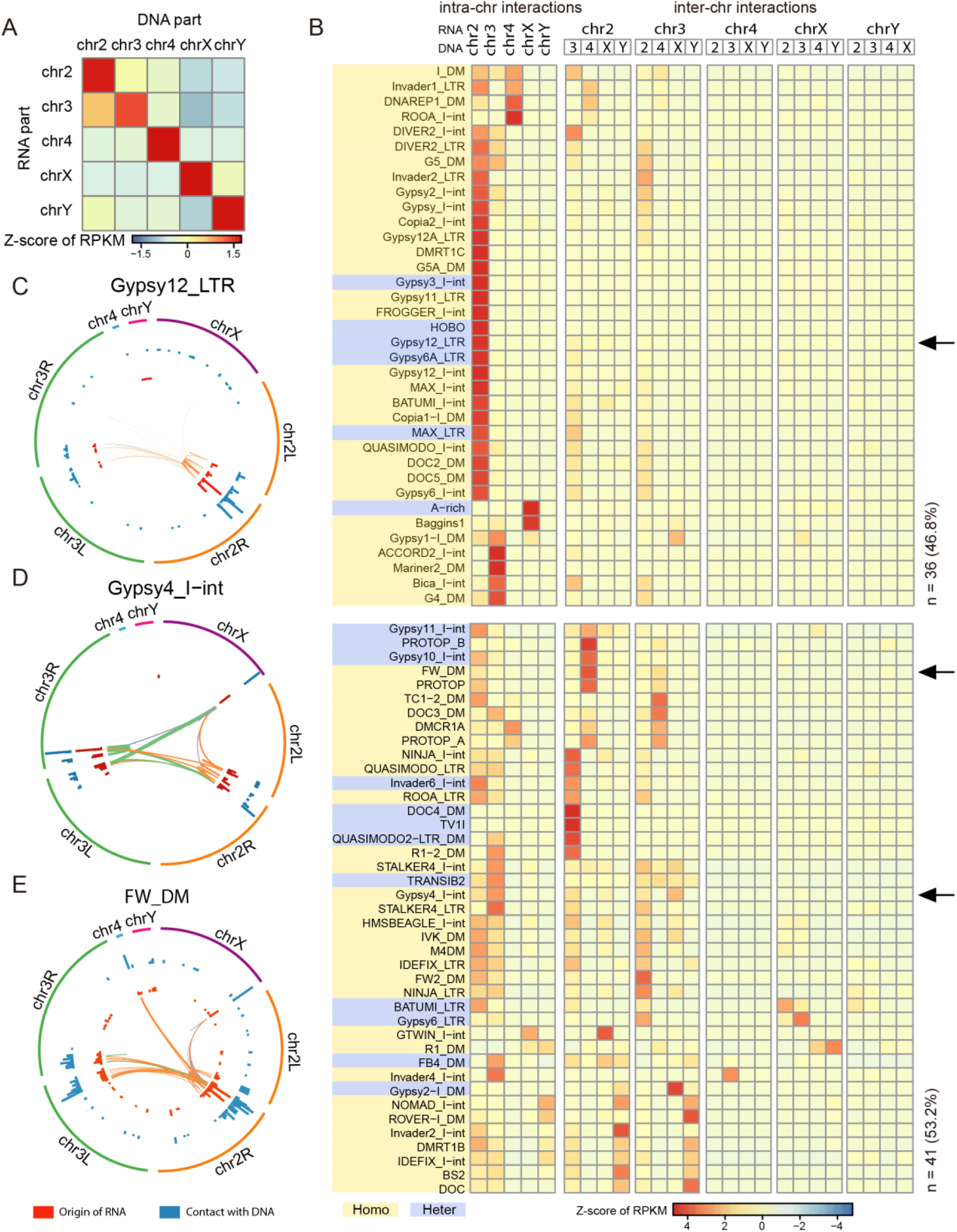
*Cis*- and *trans*-acting repeat-derived RNAs on chromatin. (A) Heatmap of RNA-DNA interaction scores for 77 CHARRs on the same or different chromosomes. *Trans*-chromosome interactions are referred to as RNA interacts with DNA in chromosomes other the RNA is transcribed from. (B) Assignment of each of the 77 CHARRs to either intra- or inter-chromosomal interactions in S2 cells and sorting of the data by unsupervised clustering. Top: RNA derived from different chromosomes (first row) and its DNA contacts in the same or different chromosomes (second row). The RNA-DNA interaction intensities were indicated by the z-scores according to the color key at the bottom. Arrows point to the three specific CHARRs Gypsy12_LTR, Gypsy4_I-int and FW_DM, as individually illustrated in panel C, D, and E. All CHARRs are either highlighted with yellow or blue to indicate their preferences for homotropic or heterotropic interactions as described later. (C, D, E) The origins of three representative CHARRs and their DNA contacts, as shown by the Circos plots for Gypsy12_LTR (C), Gypsy4_I-int (D) and FW_DM (E). In each of these plots, the inner (red) track indicates the origins of individual RNAs and the outer (blue) track shows where the RNAs interact with DNA in 100kb DNA windows. The heights of the signals correspond to Reads per 100kb window per millions. Lines specify intra- or inter-chromosomal interactions with width indicating the relative interaction levels in each case. The line color indicates the origin of chromatin-interacting RNAs.

### Tendency for *trans*-acting RNAs to supplement *cis*-acting RNAs on chromatin

During our analysis of *cis*- and *trans*-acting CHARRs, we further noted an interesting phenomenon where DNA loci expressing high levels of CHARRs tended to associate with less *trans*-acting RNAs (related repeat-derived RNA species transcribed from other chromosomes) and the converse was also true, as exemplified on a specific region in chromosome 2R where TV1I RNA (a Gypsy family member from the LTR class) was mainly transcribed from region 1 (Fig. 5A, second track), most of which contacted DNA locally around the transcribing locus (Fig. 5A, third track). Interestingly, *trans*-acting RNAs (transcribed from TV1I-related repeat species from other chromosomes) were mostly mapped to region 2 where little TV1I RNA was produced and 3 where a much lower level of the TV1I RNA was transcribed compared to region 1 (Fig. 5A, third track). We quantified these results by dividing all active TV1I loci into 3 groups according to their levels of transcription (bottom 25%, middle 50%, and top 25%, Fig. 5B), and then determined the ratio of *trans*-acting over *cis*-plus *trans*-acting RNAs for each group and observed a reverse correlation between local RNA production and the percentage of association with *trans*-acting RNAs (bars, Fig. 5B). This phenomenon also applied to another repeat RNA, Baggins1 (a LOA family member from the LINE class) (Fig. 5C, D). To determine whether this represents a general trend, we extended the analysis to all CHARRs and found that, with a few exceptions, most followed this rule (Fig. 5E). Together, these observations suggest that highly transcribed CHARRs supply RNAs in *trans* to interact with DNA regions with less transcription activities.

**Figure 5.**
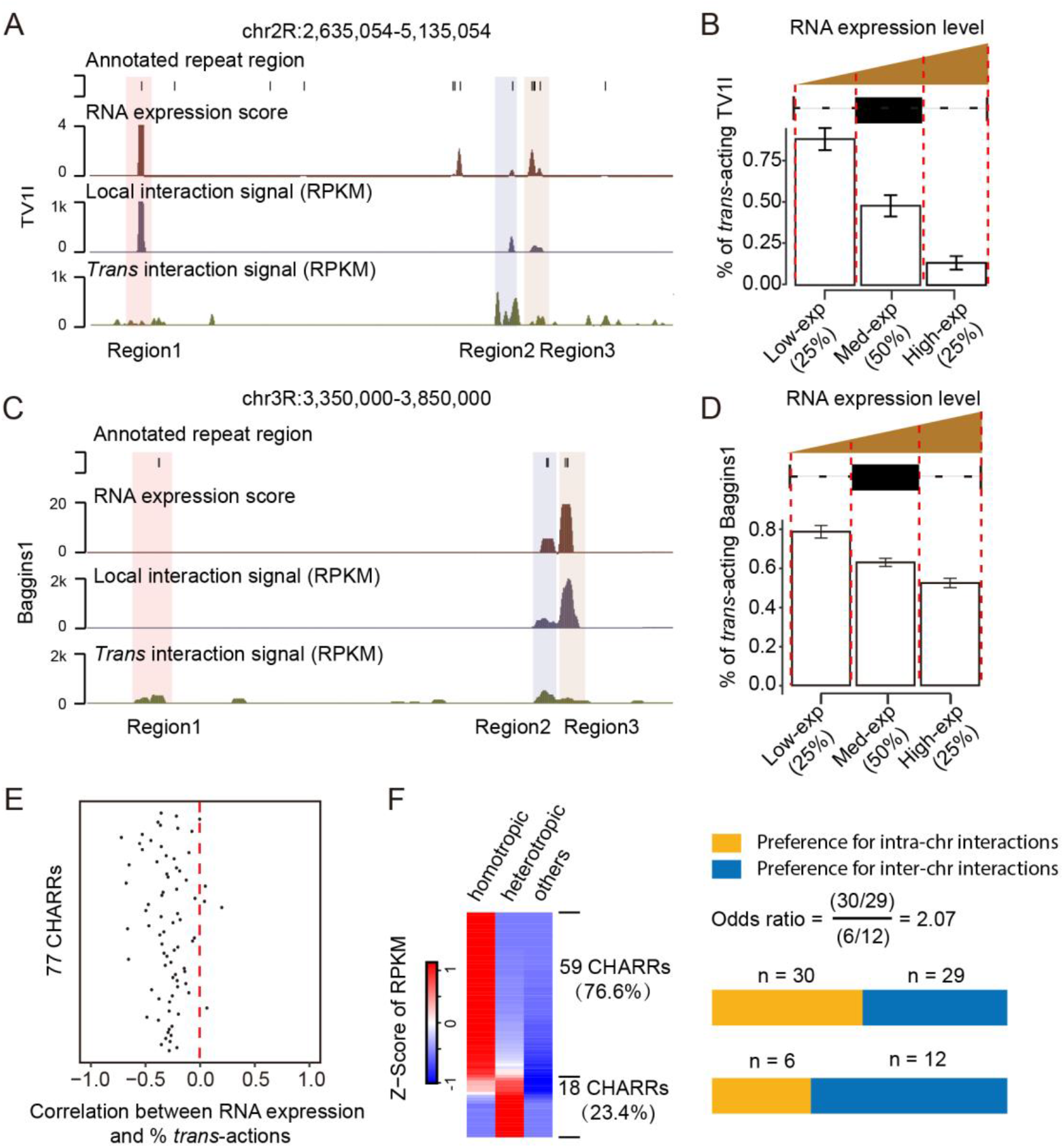
Reverse correlation between local transcription and association with *trans*-acting RNAs on chromatin. (A) A representative genomic region showing TV1I transcription loci and interactions with TV1I subfamily RNAs produced either locally or from other chromosomes (*trans*-acting RNAs). Three annotated TV1I transcription regions are indicated at bottom. (B) TV1I subfamily-derived RNAs were segregated into three groups according to their levels of transcription (bottom 25%, middle 50%, and top 25%). Bars indicate the percentage of *trans*-acting TV1I RNAs on the transcription loci in each group. (C and D) Similar analysis and illustration for another typical CHARR Baggins 1 as in A and B.(E) Pearson correlation scores for individual CHARRs between their expression and the percentage of associated *trans*-acting RNAs from the same subfamilies.(F) Tendencies of CHARRs engaging homotropic versus heterotropic interactions based on normalized reads per kilobase per million (Left). The two classes of CHARRs were further separated into those with preference for inter- (yellow) or intra- (blue) chromosomal interactions (Right), as indicated in Fig. 4.

Given the observation that some CHARRs preferred to interact with DNA near their sites of transcription while others showed both *cis*- and *trans*-chromosomal interactions, we next characterized the underlying DNA sequences that might specify such RNA-DNA interactions. If a given CHARR specifically interacted with a DNA region that harbors the same repeat sequence within a 1kb window, we called it “homotropic” interactions. If the interacting DNA region contains distinct repeat sequence (e.g. those encoding for RNAs of different classes or different subfamily members), we then referred it to as “heterotropic” interactions. All DNA interaction regions that contain no repeat sequence were classified as “others”. According to these definitions, we found that 59 CHARRs (yellow-labeled in Fig. 4B) were predominantly engaged in homotropic interactions, whereas the remaining 18 (blue-labeled in Fig. 4B) were more involved in heterotropic interactions, and none showed significant interactions with DNA that contain no repeat sequence (Fig. 5F, left). We further noted that relative to CHARRs with preference for homotropic interactions, more CHARRs with higher tendency to engage in heterotropic interactions were more involved in inter-chromosomal interactions, as indicated by a significant odds ratio (Fig. 5F, right). These observations imply that CHARRs with preference for homotropic interactions may facilitate initiating heterochromatin formation, whereas those for heterotropic interactions may play more important roles in heterochromatin spreading as well as maintenance.

### Dicer-2 processed repeat-derived RNAs for heterochromatin maintenance

Most retrotransposons are known to be bi-directionally transcribed. To verify this, we took advantage of the existing GRO-seq data on *Drosophila* S2 cells (Supplementary Table 2), showing that both sense and anti-sense repeat-derived transcripts were indeed represented in relatively equal abundance, with rRNAs served as control for predominantly sense transcription (Supplementary Fig. 6A). Interestingly, but not necessarily surprisingly, the identified CHARRs were among the most abundant repeat-derived RNAs. This was true regardless of different RNA classes (Supplementary Fig. 6B), which is in contrast to annotated non-repeat transcripts, the majority of which was transcribed from the sense strand (Supplementary Fig. 6C), with exception of some long intergenic non-coding RNAs (Supplementary Fig. 6D).

Bi-directionally transcribed repeat RNAs may thus provide dsRNA substrates for further processing into endo-siRNAs to function in heterochromatin formation/maintenance, as demonstrated earlier(Fagegaltier et al. 2009; Volpe and Martienssen 2011). Furthermore, in contrast to piRNA-mediated heterochromatin formation in germline, Dicer-2 has been reported to be specifically devoted to endo-siRNA processing in fly somatic cells(Lee et al. 2004; Pham et al. 2004; Czech et al. 2008; Ghildiyal et al. 2008). To determine whether the CHARRs we identified all depended on Dicer-2 for their efficient processing and thus expression, we took advantage of the existing small RNA-seq data (Supplementary Table 2) to compare their expression levels between wild-type, Dicer-2 knockout, and Dicer-2 rescued S2 cells(Kandasamy and Fukunaga 2016). We found that almost all CHARRs were down regulated in response to Dicer-2 knockout and rescued in Dicer-2 re-expressed cells (Supplementary Fig. 7A). Note that much higher expression of those endo-siRNAs in Dicer-2 re-expressed cells likely resulted from Dicer-2 overexpression. Importantly, the RNA-seq reads of total CHARRs from wild-type S2 cells were distributed between 18 to 25nt in length, consistent with their processing into endo-siRNAs (Supplementary Fig. 7B). These endo-siRNAs appeared to have assembled into Ago2-containing complexes, as 54 (75%) CHARRs could be identified in the published Ago2 RIP data (Supplementary Fig. 7C). Therefore, re-analysis of these published data on CHARRs strongly suggests that Dicer-2 is responsible for processing CHARRs into endo-siRNAs to help maintain heterochromatin in S2 cells.

To directly test this hypothesis, we performed Dicer-2 knockdown in S2 cells (Supplementary Fig. 7D, E) and confirmed drastic reduction of small RNAs derived from DMCR1A or DOC by Northern blotting (Fig. 6A). Because many CHARRs were able to supply RNAs in *trans*, as we documented in the present study, we asked whether or not the associated phenotype previously detected on heterochromatin could be “rescued” with small RNAs derived from CHARRs. For this purpose, we chemically synthesized a pool of endo-siRNA mimics based on a representative subset of CHARRs (DMCR1A, FB4_DM, FB_DM, Gypsy4_I-inU, DOC, Gypsy2-I and I_DM, Supplementary Table 3) and transfected this pool into S2 cells depleted of Dicer-2. We found that relative to wild-type cells, Dicer-2 knockdown reduced H3K9me3 as expected, and importantly, the CHARR-derived siRNA pool, but not scrambled siRNA, effectively restored this heterochromatin marker, while the levels of H3K27me3 and H3K4me3 remained constant under these experimental conditions (Fig. 6B, quantified below based on triplicated experiments). This was also evident at the immunocytochemical level by staining for H3K9me3 (Fig. 6C). These data suggest that the CHARR-derived siRNA mimics were able to rescue heterochromatin defects upon transfection (thus only acting in *trans*) into Dicer-2 deficient S2 cells.

**Figure 6.**
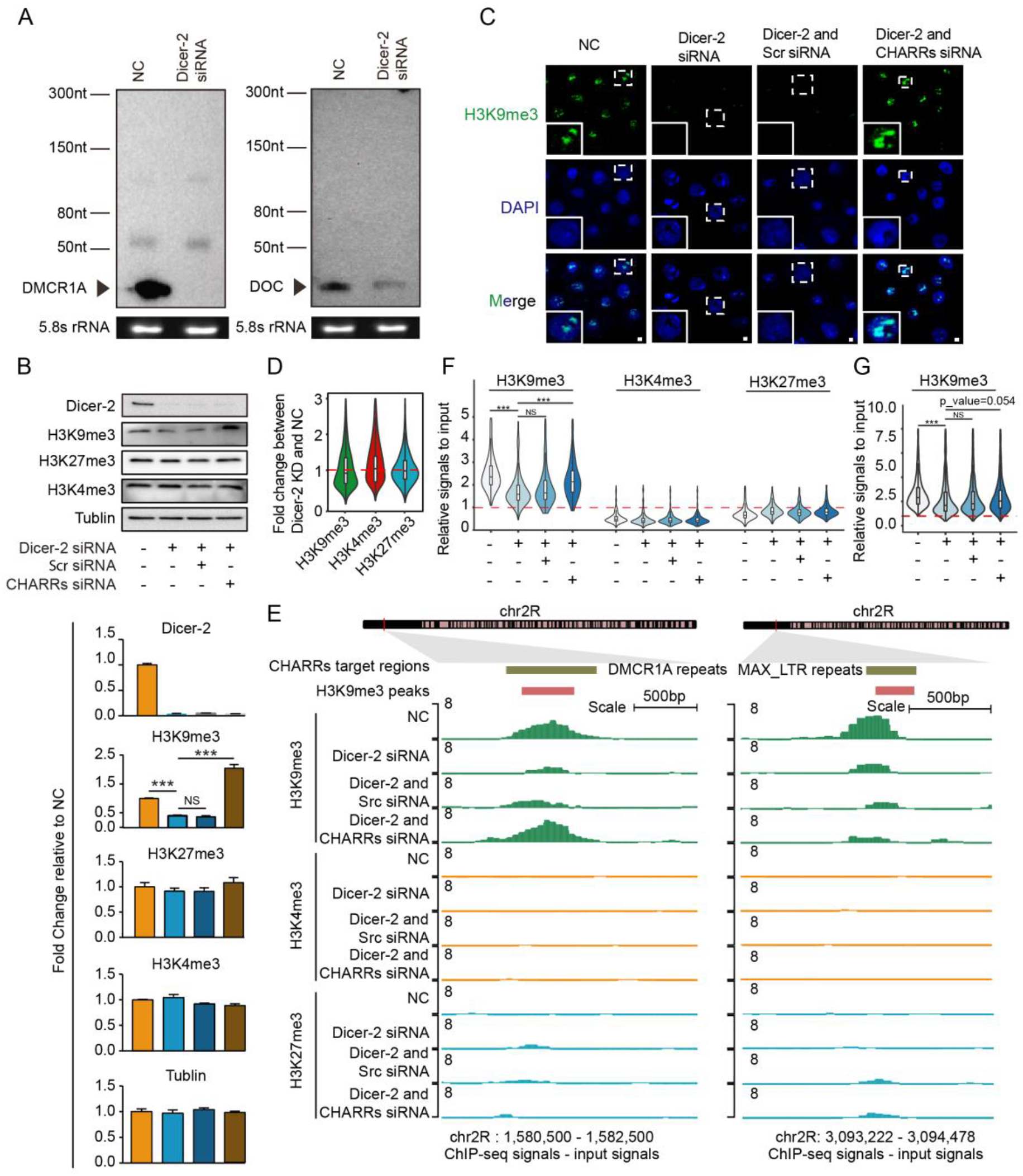
Trans-acting repeat-derived RNAs for heterochromatin maintenance. (A) Confirmation of Dicer-2 dependent expression of DMCR1A (left) and DOC (right) expression by Northern blotting analysis.(B) Western blotting analysis of Dicer-2, H3K9me3, H3K27me3, H3K4me3 and Tubulin in S2 cells in response to siRNA-mediated knockdown of Dicer-2 and rescue with a transfected pool of CHARR-derived synthetic siRNAs. Quantified data are shown below as fold-change (FC) relative to input (NC) in lane 1. Data are presented as mean ± SEM (n = 3 biological replicates). *p<0.05, **p< 0.01, ***p<0.001. (unpaired Student’s t test).(C) H3K9me3 detected by immunocytochemistry in S2 cells treated with different combinations of siRNAs, as indicated. Green, H3K9me3 signals; blue, DAPI. Scale bar, 2μm.(D) Violin plot for fold-change of H3K9me3, H3K4me3 and H3K27me3 signals on CHARRs-targeted peaks in response to Dicer-2 knockdown. KD: knockdown. (E) H3K9me3 ChIP-seq signals on representative genomic loci, one corresponding to a CHARR-target locus (left) and a non-CHARR-target locus (right) in response to Dicer-2 knockdown, complemented with either scrambled or CHARRs-derived siRNAs. ChIP-seq signals for euchromatin (H3K4me3) or facultative heterochromatin (H3K27me3) were shown for comparison.

### Rescuing global heterochromatin defects in Dicer-2 deficient cells

To demonstrate the contribution to repeat-derived endo-siRNAs to heterochromatin maintenance genome-wide, we next performed ChIP-seq for H3K9me3 in comparison with H3K4me3 and K3K27me3 in response to Dicer-2 knockdown with or without treatment with CHARR-derived siRNA mimics in S2 cells. Our ChIP-seq datasets were comparable to the profiles of these histone marks published earlier on the same cell type, demonstrating the quality of our data (Supplementary Fig. 8A, B). Upon Dicer-2 knockdown, we detected global decrease of the H3K9me3 ChIP-seq signals, while the ChIP-seq signals for H3K4me3 and H3K27me3 were not affected (Fig. 6D). The effects were also evident on individual H3K9me3-marked genomic loci (see first two tracks in Fig. 6E), suggesting that Dicer-2 is functionally required for maintaining heterochromatin genome-wide.

The next question was whether CHARR-derived siRNA mimics were able to rescue heterochromatin defects, and if so, whether the rescue required their targeting specificity. Because those siRNA mimics were designed to target 10 representative CHARRs (Supplementary Table 3), we thus analyzed the H3K9me3 ChIP-seq signals on 1kb-binned genomic regions that show homology with at least one of the CHARR-derived siRNA mimics (Fig. 6F) in comparison with genomic regions that showed H3K9me3 ChIP-seq signals but with <50% homology with any of those CHARR-derived siRNA mimics (Fig. 6G). Indeed, we found that CHARR-derived siRNA mimics, but not scrambled siRNA, effectively rescued H3K9me3 ChIP-seq signals on CHARRs target genomic regions (compare lane 2 vs. 4 in Fig. 6F), but modest at best (likely due to a remaining degree of heterotropic interactions) on none CHARRs target regions (compare lane 2 vs. 4, p=0.054, in Fig. 6G). As expected, little H3K4me3 and H3K27me3 ChIP-seq signals were detected in H3K9me3-marked genomic loci. These general trends were also illustrated on two representative genome loci for CHARRs targets (Fig. 6E, left and Supplementary Fig. 9A) and none CHARRs targets (Fig. 6E, right and Supplementary Fig. 9B). Collectively, these data demonstrated that a pool of *trans*-acting CHARR-derived siRNA mimics was able to bypass the functional requirement of Dicer-2 for maintaining heterochromatin homeostasis on their target regions in *Drosophila* S2 cells.

### CHARR-derived endo-siRNAs required for faithful chromosome segregation

It has been previously documented that knockdown of Dicer-2 and Ago-2 caused significant defects in chromosome segregation during cell division in S2 cells(Pek and Kai 2011). Because Dicer-2 and Ago-2 are key siRNA pathway components, it is reasonable to extrapolate that endo-siRNAs processed by Dicer-2 and loaded on Ago-2 are responsible for the phenotype, but direct evidence for this critical conclusion has been lacking. Given that the CHARRs we identified cover most of pericentromeric regions to maintain the heterochromatin status in the fly genome (see Fig. 3B) and most of these CHARRs-derived were bound by Ago-2 (see Supplementary Fig. 7C), we took advantage of the ability of CHARRs-derived siRNAs to rescue most heterochromatin defects to ask whether these siRNA mimics were also able to rescue the chromosome segregation defects. To this end, we first confirmed that knockdown of either Dicer-2 or Ago-2 caused cell cycle defects, and as expected, we detected G1-S arrest in both cases (Fig. 7A, B and Supplementary Fig. 10). Importantly, we found that CHARRs-derived siRNAs, but not scrambled siRNA, were able to rescue the cell cycle defects in Dicer-2 knockdown cells (Fig. 7A), but not Ago-2 knockdown cells (Fig. 7B). These data are fully in line with the requirement of Dicer-2 for processing CHARRs into small endo-siRNAs, which could be bypassed by the transfected siRNA mimics, although these siRNA mimics would still need Ago-2 to execute their functions in the cell.

**Figure 7.**
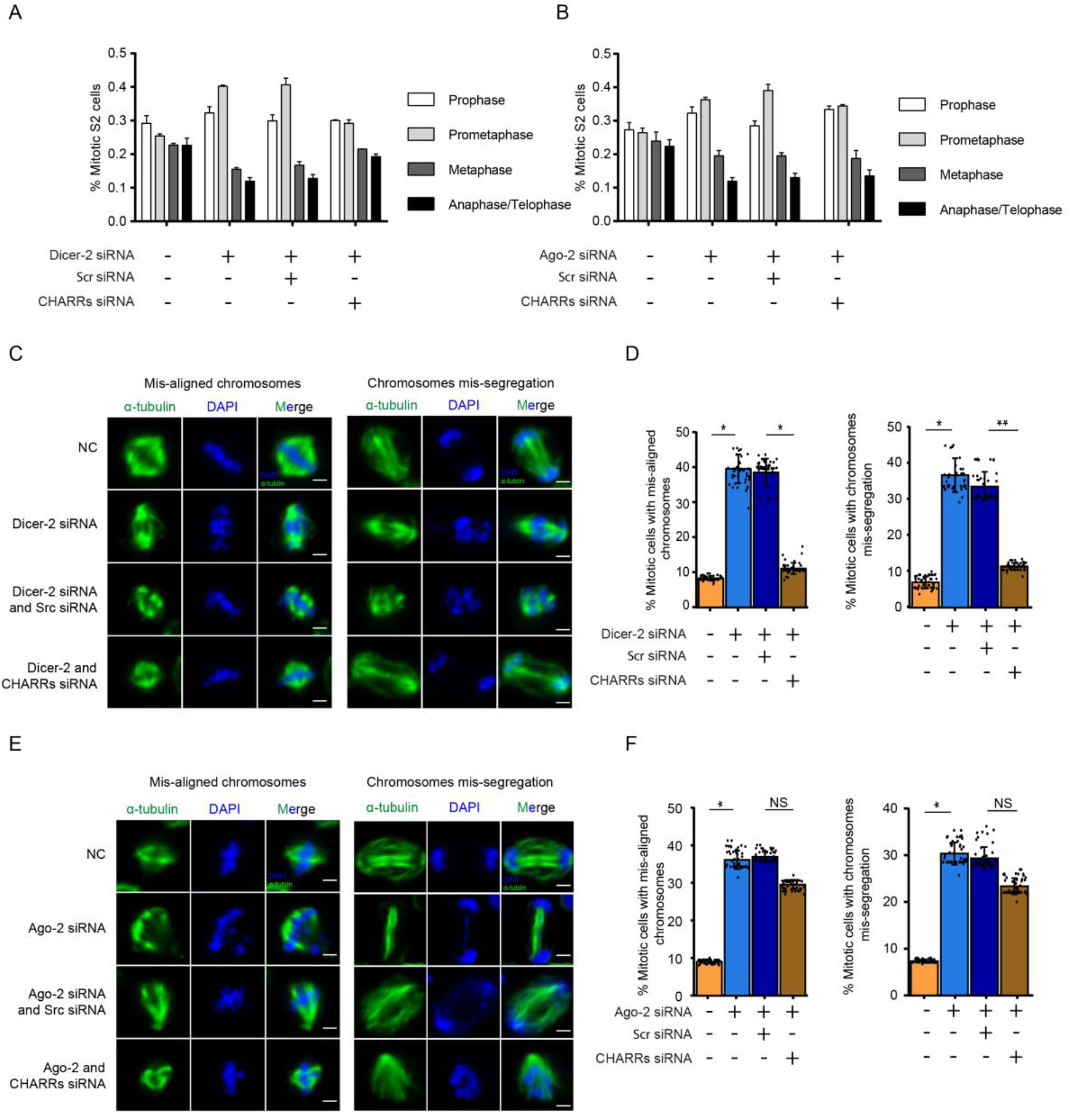
CHARRs could rescue Dicer-2 knockdown, but not Ago-2 knockdown, induced cell division defects. (A and B) Percentages of S2 cells at each stage of mitosis in response to knockdown of Dicer-2 (A) or Ago-2 (B) and treatment with either scrambled or CHARRs-derived siRNAs. n=50 for each experiment. (C) Dicer-2 knockdown-induced chromosomes mis-alignment (left) and mis-segregation (right) and rescue by CHARRs-derived siRNAs, but not scrambled siRNA. Green, stained α-tubulin; blue, DAPI. Scale bar, 2μm.(D) Percentages of S2 cells at metaphase exhibiting mis-aligned chromosomes (left) and anaphase exhibiting lagging chromosomes (right) at different experimental conditions. n=50 for each condition. *p<0.05, **p<0.01, NS: not significant (multiple group Student’s t test). (E and F) Similar to C and D except on Ago-2 knockdown cells. n=50 for each condition. *p<0.05, **p<0.01, NS: not significant (multiple group Student’s t test).

We closely examined the mitotic defects in Dicer-2 knockdown cells, noting both mis-aligned chromosomes and lagged as well as mis-segregated chromosomes (Fig. 7C and Supplementary Fig. 11A) in about equal frequencies (Fig. 7D). Importantly, transfection of CHARRs-derived siRNA mimics into these Dicer-2 deficient cells, but not scrambled siRNA, were sufficient to rescue these defects (Fig. 7D and Supplementary Fig. 11A). In contrast, we observed similar mitotic defects in Ago-2 knockdown cells, but the CHARRs-derived siRNAs failed to correct the phenotype (Fig. 7E and Supplementary Fig. 11B). Taken together, these results provide unequivocal evidence for active retrotransposon RNAs to preserve pericentromeric heterochromatin homeostasis and thus cell cycle progression through the Dicer-2/Ago-2 mediated endo-siRNA pathway.

## DISCUSSION

Genetic and biochemical experiments in fission and *Drosophila* have laid a general conceptual framework in understanding the formation and maintenance of constitutive heterochromatin in centromeric and pericentromeric regions(Ekwall et al. 1995; Kellum and Alberts 1995; Grewal and Jia 2007). Interestingly, however, while heterochromatin is in general prohibitive to transcription, mounting evidence suggests that transcription is actually required to initiate heterochromatin formation. This so-called nascent RNA model(Buhler et al. 2006; Holoch and Moazed 2015a) also creates a puzzle in envisioning how heterochromatin could be actively maintained. As a matter of fact, it has been reported that repeat-derived RNAs are prevalent on chromatin in vertebrate cells from *Drosophila* to humans(Hall et al. 2014), implying that there is no shortage of repeat-derived RNAs to help maintain heterochromatin. The central question is what is the nature of these RNAs and where they come from?

The consensus in the field is that heterochromatin-associated RNAs are mostly derived from active retrotransposons and simply repeats. However, the nature of such repetitive sequences has made it difficult to determine their origins and destinations in the genome. We have now attacked this fundamental problem by using the newly elucidated RNA-DNA interactome in *Drosophila* somatic cells(Li et al. 2017). Taking advantage of such high density interactome, we now show that various active retrotransposons, especially those from the Gypsy family of the LTR class, produce a large amount of repeat RNAs that are selectively associated with pericentromeric regions in the fly genome. Interestingly, while a subset of those repeat-derived RNAs tends to act in *cis*, another subset appears to interact with DNA in both *cis* and *trans* modes. Particularly interesting is a general trend we have uncovered, where DNA loci with more active local transcription seem to inversely correlate to their ability to attract *trans*-acting RNAs from related repeat species. Therefore, RNAs released from those more active loci may supply extra RNAs to act in *trans* on less transcribed loci. This “community” act of repeat-derived RNAs may thus help “patch up” certain heterochromatic regions “damaged” by chromatin remodeling activities in the cell, such as that catalyzed by the H3K9me3 demethylase, thereby ensuring heterochromatin homeostasis for stable epigenetic inheritance.

In fission yeast, repeat-derived RNAs are amplified by an RNA-dependent RNA polymerase, which may supply a population of *trans*-acting RNAs to help initiate and/or maintain heterochromatin in other genomic loci(Yu et al. 2018). A different mechanism is employed in *Drosophila* germline where repeat-rich transcripts are processed and amplified by the piRNA machinery to ensure efficient silencing of retrotransposons to protect the genome integrity as well as to help maintain heterochromatin in centromeric and pericentromeric regions to ensure accurate chromosome segregation during cell division in gonad(Muerdter et al. 2013; Iwasaki et al. 2015). The problem is that such RNA amplification mechanism does not seem to exist in somatic cells of flies and mammals. The endo-siRNA pathway has clearly been implicated in *Drosophila* somatic cells(Czech et al. 2008; Ghildiyal et al. 2008), although it remains unclear whether such endo-siRNA pathway also operates in somatic cells of mammals, which might result from less attention paid on processing of repeat-derived RNAs by the conserved siRNA machinery known to be highly active in mammals. In any case, at least in *Drosophila* S2 cells, we now document that such endo-siRNA pathway is responsible for generating small RNAs that can act in *cis* and *trans* to help maintain heterochromatin homeostasis in pericentromeric regions. We envision a similar mechanism that may also operate in somatic cells in mammals, which requires future investigation once the RNA-DNA interactome of much higher density becomes available.

Knockdown of Dicer (Dicer-1 in mammals and Dicer-2 in fly) and Ago-2 has been shown to cause mitotic defects in both fly and mammalian cells(Pek and Kai 2011; Huang et al. 2015), but it has been unclear whether these small RNA machineries act through their traditional functions or through some new mechanism(s) in the nucleus. We now show that a pool of synthetic repeat-derived siRNA mimics is able to rescue all measurable cell cycle defects in *Drosophila* S2 cells, which is fully compatible to the central role of the endo-siRNA pathway in ensuring cell cycle progression through maintaining pericentromeric heterochromatin. We speculate the conversed function of this pathway in mammals, although different classes of retrotransposons are likely involved in different organisms.

The production of repeat-derived small RNAs to help maintain heterochromatin for stable genetic inheritance in somatic cells suggests a key and immediate benefit of active retrotransposons for the genome. This realization is interesting because retrotransposons have been traditionally viewed as mutagens in the genome, although their relatively random actions may facilitate genome evolution in the long run. In *Drosophila* germ cells, an RNA amplification mechanism has been evolved to maximally suppress this mutagen function of retrotransposons to ensure genome integrity. We now show that despite the lack of such RNA amplification mechanism in somatic cells, the endo-siRNA machinery is still quite active to ensure the supply of repeat-derived small RNAs. Therefore, without worrying about the responsibility to transmit genetic materials to offspring, somatic cells may tolerate potential genome vulnerability in exchange for stable genetic inheritance during development and differentiation. As higher eukaryotic genomes are populated with enormous amounts of repeat sequences, we suggest that some of those “junk” DNA sequences actually have important functions while most others are fossil of genome evolution.

## METHODS

### Cell Lines and Cell Culture Conditions

S2 cells were cultured under sterile conditions at 26°C in Schneider medium (Invitrogen) containing 10% heat-inactivated fetal bovine serum and 100 µg/ml penicillin-streptomycin. S2 cell lines are negative for mycoplasma contamination.

### Alignment of GRID-seq reads to the *Drosophila* genome

GRID-seq raw reads were split into RNA and DNA reads according to the designed bivalent linker. Trimmomatic(Bolger et al. 2014) was used to remove adapter sequences and filter low-quality reads by using the parameters *MINLEN: 18* and *SLIDINGWINDOW: 2:20*. Filtered reads were aligned to the *Drosophila* genome (genome version :dm6) with ShortStack(Axtell 2013) using the parameter *–m 400* for DNA reads and *–m 200* for RNA reads, respectively. Multiple mapped reads were weighted based on the frequencies of neighbor uniquely mapped reads (Supplementary Fig. 1C). Unmapped reads were cleaned with SAMtools(Li et al. 2009) using the parameter *–F 4*, and the weighted score of each read was recorded in the fifth column of bed file.

### Annotation of RNA and DNA reads

Annotated unique RNAs of *Drosophila* was downloaded from FlyBase(Drysdale 2008) and the repeat sequences from RepeatMasker track in the UCSC genome browser(Jurka et al. 2005). RNA reads were annotated to unique and repeat RNAs using IntersectBed tool with the parameter -*f 1.0, -split* and *–s.* The *Drosophila* genome was scanned with EMBOSS (Rice et al. 2000) based on Alu I restriction sites from REBASE (Roberts et al. 2015) to generate AluI DNA bins. IntersectBed tool was used to connect Alu I DNA bins using the parameter *-f 1.0*.

### Assigning RNA-DNA interactions

Multiple read pairs that have the same mapped RNA and DNA loci associated with the same PCR primer sequences were considered PCR duplicated, and thus were counted only once. The following equation was used to compute RNA-DNA interactions in each AluI DNA bin:

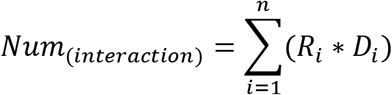

Where *n* is the number of assigned read pairs. For each read pair *i*, *R*_*i*_ is the contribution score of this read to the RNA part and *D*_*i*_ is the contribution score of this read to the interacting AluI DNA bin. *R*_*i*_=1, if the RNA part of the read pair is uniquely mapped, or equals to a fraction based on the weighted score.

### Construction of non-specific background using mixed GRID-seq libraries

A mixed GRID-seq library from human MDA-MB-231 and *Drosophila* S2 cells was used to construct the non-specific RNA-DNA interaction profile. RNA reads, which were only mapped to the human genome (hg38) using Bowtie(Langmead et al. 2009) with the parameter *–n 0*, were kept. Their mated DNA reads were processed using ShortStack, as described above. The human RNA signals within each 1kb DNA bin were normalized to one million, which was further smoothed by a moving window that includes 5 upstream and 5 downstream bins. The final coverage of the 1kb DNA bin *i* is:

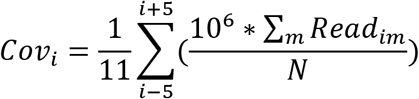

where *m* is the number of reads mapped to the1k DNA bin *i* and *N* is the total read number mapped to the *Drosophila* genome.

To make DNA binding scores comparable between Alu I binned versus 1kb binned genome, RNA binding signals in each 1k DNA bin were converted to RNA binding signals in each AluI DNA bin by first dividing each 1kb DNA bin into 1000 1bp bins to calculate the signal in each small bin based on 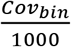. IntersectBed tool with the parameter *–f 1.0* was used to compute signals in each Alu I DNA bin by summing the signals from all 1bp bins in the fragment. Finally, signals in AluI DNA bins were all normalized to signals per 1kb:

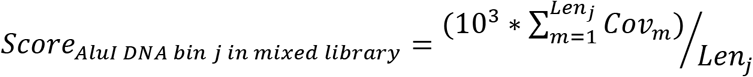

where *Len*_*j*_ is the length of the AluI DNA bin *j*.

### Filtering singular and background to identify specific RNA-DNA interactions

To support specific RNA-DNA interactions, we required at least two RNA-DNA mates for each RNA transcript in a given Alu I DNA bin. We also developed two background models to simulate Poisson distribution of RNA binding signals on DNA. The first was based on uniform distribution of individual RNAs on DNA, based on which we estimated the background score:

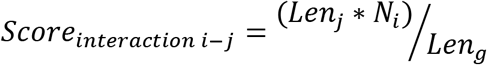

where the *Len*_*g*_ is the length of the genome, *Len*_*j*_ is the length of AluI DNA bin *j* and *N*_*i*_ is total signals for RNA *i*. This score was further normalized according to length in each AluI DNA bin, and the resulting RPK (reads per kilobase) was used as the *λ*_*BG*_*i*_ value to obtain the Poisson distribution of this RNA-engaged genomic interactions and to calculate the p-value for such interactions. The ratio of *Num*_*interaction i*−*j*_ over *Score*_*interaction i*−*j*_ was reported as the fold_change (FC) above the background. We also developed a second background model with that data deduced with RNA signals from the mixed library. We first calculated the non-specific interaction score (*Score*_*AluI DNA bin j in mix library*_) based on human-derived RNA binding signals, and then used this score as the *λ*_*BG*_*mix*_ value to obtain the Poisson distribution of human RNA-engaged genomic interactions and to calculate the p-value for such interactions. The ratio of length and sequencing depth normalized *Num*_*interaction i*− *j*_ (in RPKM) over *λ*_*BG*_*mix*_ was reported as the fold_change (FC). RNA-DNA interactions that met the requirement of p-value <0.05 and the fold_change (FC) >2 based on both background models were considered specific and thus retained for further analysis.

### Data normalization for comparison between GRID-seq libraries and different RNAs within the same libraries

The interaction RNA-DNA score for each RNA is affected by the sequencing depth in different GRID-seq libraries, the length of each Alu I DNA bin, and the length of each RNA. To enable comparison among different libraries and different RNAs within the same libraries, we normalized these variables according to:

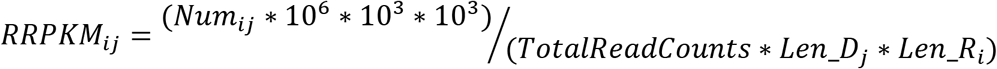

where *Num*_*ij*_ is the interaction score of RNA *i* with Alu I DNA bin *j*, *TotalReadCounts* is the sequencing depth of individual libraries, *Len*_*D*_*j*_ is the length of Alu I DNA bin *j* and *Len*_*R*_*i*_ is the length of RNA *i*.

### Comparison between GRID-seq and ChAR-seq datasets

ChAR-seq raw data were downloaded from the GEO database (Supplementary Table 2). PCR duplicates were removed using Clumpify with default parameters. All five independent ChAR-seq libraries were combined followed by adapter trimming and filtering low-quality reads with Trimmomatic using the parameters *MINLEN: 36, LEADING:3 TRAILING:3* and *SLIDINGWINDOW: 4:15*. Filtered ChAR-Seq reads were split into paired RNA and DNA reads, and then processed as described above for the GRID-seq data. FeatureCounts was used to calculate reads in 1kb DNA bins and then converted to RPKM using edgeR package in R language. For validation for assigning multi-mapped reads, we only considered 1kb DNA bins that contain newly assigned multi-mapped reads from the GRID-seq dataset.

### Comparison of RNA-DNA interactions in relationship with chromatin marks

Public ChIP-seq data from S2 cells were downloaded from the SRA database (Supplementary Table 2). Fastq-dump were used to covert raw SRA data to Fastq. Quality control and data processing were similar to the procedures for processing the GRID-seq data with parameters adjusted to *MINLEN: 36* and *SLIDINGWINDOW: 4:20* according to the read length. Filtered reads were mapped to the reference *Drosophila* genome (dm6) using STAR (Dobin et al. 2013) with the parameters: *--outFilterScoreMinOverLread 0.1*, *–outFilterMatchNminOverLread 0.1*, *-- alignIntronMax 1* and *--alignEndsType EndToEnd*. The wig files of each chromatin mark ChIP-seq dataset was obtained through STAR using the parameters: *–outWigTypewiggle read1_5p*, *-- outWigStrandUnstranded* and *--outWigNormRPM*. Enriched peaks were detected by MACS2 with input data as control. Top 500 peaks were used for further analysis.

### Identification of repeat RNA on constitutive heterochromatin

The ChIP-seq data for constitutive heterochromatin markers (H3K9me3 and HP1) were obtained from the GEO database (Supplementary Table 2) and the RPKM values were calculated on 10kb DNA bins. The Pearson correlation score was calculated on each DNA bin containing binding signals from repeat RNAs. Repeat sequences, except rRNAs, were defined as CHARRs if the correlation score with H3K9me3 or HP1 is >0.3.

### Analysis of CHARR-DNA interactions relative to Hi-C defined compartments

Public Hi-C data from S2 cells were downloaded (Supplementary Table 2) and processed with Trimmomatic using default settings plus trim tool in Homer using the parameter *-3 AAGCTT*. Trimmed reads with length of ≥38nt were mapped to the *Drosophila* genome using end-to-end alignment model provided by Bowtie2 (Ref (Langmead and Salzberg 2012)). We discarded potential PCR duplicates as well as reads with no useful information, including (1) read pairs separated <1.5× of the sequenced insert fragment length, (2) reads from 10kb regions containing >5× of the average coverage, (3) read pairs lacking restriction sites at the 3’ end of either read within the estimated fragment length, (4) reads with their ends resulting from self-ligation with adjacent restriction fragments. We then used the filtered dataset to perform PCA analysis with HOMER, using runHiCpca.pl with the parameters *–res 10kb* and *–superRes 20kb* and compartments analysis using runHiCpca.pl and findHiCCompartments.pl. Finally, CHARR-DNA interactions signals and Hi-C compartments were intersected with IntersectBed from BedTools. The length of Hi-C compartment A and B was normalized to 1Mb.

### Processing GRO-seq, small RNA-seq and Ago2 RIP data from S2 cells

All data were downloaded from public databases (Supplementary Table 2). GRO-seq data were processed as with the ChIP-seq data for chromatin marks. Annotated unique and repeat RNA species were used to calculate individual transcription scores by using FeatureCounts and EdgeR.

Small RNA-seq data were similarly processed as above using adjusted Trimmomatic parameters: *SLIDINGWINDOW:4:24* and *MINLEN:18*. SortMeRNA(Kopylova et al. 2012) were used to filter ribosomal RNAs with default setting. We also used ShortStack to map both uniquely and multi-mapped reads to obtain the RPM value for each CHARR.

Reads for the downloaded Ago2 RIP data were blasted to identify repeat RNA-derived sequences using the parameters *-outfmt 6* and *-word_size 7*. Reads will be kept for further analysis if the mismatch number is under 3 and the mapping region of the reads equals to the whole length of the reads. We then counted the reads number for each repeat sequence.

### siRNAs-mediated knockdown of Dicer-2 and Ago-2 and immunofluorescence

Transfection was performed, as described(Rogers and Rogers 2008), using the “bathing” method for siRNA delivery. Cells were counted, pelleted and resuspend at 1-5×10^6^ cells/ml in serum free media. We added ∼10-30 µg dsRNA to each well in the 6-well tissue culture plate to obtain the final concentration of 25-50 nM. About 1 ml of cells were seeded each well in the 6-well plate and incubate at room temperature for 30 min followed by the addition of 3 ml complete media with 10% FBS to each well. This process was repeated every other day for three times before harvesting the cells for downstream assays. Sequences of individual siRNAs were listed in Supplementary Table 3.

For immunostaining, S2 cells were washed in 1× PBS and fixed in 4% paraformaldehyde (pH-7.2) for 10 min at room temperature. After washing four times, cells were permeabilized with 0.1% Triton X100 in 1× PBS for 5 min at room temperature. Permeabilized cells were then incubated in 1% normal goat serum in 1× PBST for 30 min at room temperature, and then with the primary antibody (α-rabbit H3K9me3 1:500) and secondary antibodies (1:400) both in blocking buffer (3% BSA, 1% goat serum in PBST), each for 1h at room temperature. Cells were washed three times at room temperature, each for 5 min with 1× PBST. After mounting on coverslip with DAPI, cells were examined under Zeiss LSM-700 confocal laser scanning microscope.

### ChIP-seq library construction and data analysis

DNA libraries were constructed using the NEBNext® Ultra™ II DNA Library Prep kit (NEB, USA) following manufacturer’s recommendations. After end repair, 5’ phosphorylation. and dA-tailing of purified DNA fragments, NEBNext adaptors with hairpin loop structure were ligated and library fragments were purified with SPRIselect sample purification beads (NEB, USA). After ensuring the quality of libraries on the Agilent Bioanalyzer 2100 system, individual libraries were sequenced on the Illumina HiSeq X Ten platform to generate 150 bp paired-end.

Adapters and low-quality reads were filtered to obtain clean reads and clean reads mapped to the reference *Drosophila* genome (dm6) using Bowtie2. The wig files of each chromatin mark ChIP-seq dataset was obtained by using bamCoverage tools from Deeptools(Ramirez et al. 2016) with the parameters*: --binSize 1000, --normalizeTo1x 142573017 and --ignoreForNormalization chrM.* Enriched peaks were detected by MACS2 with input data as control using the parameters *-f BAMPE --nomodel --keep-dup all --broad*.

### Analysis of mitotic defects

Medium and fetal bovine serum were batch tested for support of normal cell growth and RNAi efficiency. For the knockdown group, specific synthetic siRNA was added to cell culture in 24-well plates. After siRNA treatment for 4 days, cells were resuspended and transferred to glass-bottom, 24-well plates (Cellvis) and allowed to adhere for 2.5 hrs before fixation, as described(Goshima et al. 2007). Cells were fixed in 4% paraformaldehyde for 10 min, permeabilized with 0.1% Triton X100 in PBS for 5 min and incubated overnight at 4°C with anti-α-tubulin (ab7291 from abcam; 1:1000) in PBS containing 0.1% Triton and 0.5 mg/ml BSA, followed by staining with secondary antibodies and DAPI (1 µg/ml). For the rescue group, siRNAi treatment was performed as above followed by transfection of CHARRs siRNA 2 days later. Immunostained specimens were imaged under a Zeiss LSM-700 confocal laser scanning microscope, using a 63x 1.4 NA oil immersion objective to achieve high resolution. We imaged two channels (DAPI, AF488) at typically 10-20 sites/well to obtain 50 metaphase cells/well on average.

### Western blotting, RT-qPCR and Northern blotting

Total cell lysate containing 15-25 μg protein from S2 cell cultures were fractionated by SDS-PAGE, immunoblotted and probed with specific antibodies. After incubation with peroxidase-conjugated secondary antibodies (1:5000; Abcam), blots were developed with Supersignal West Pico Chemiluminescent Substrate (Pierce) and exposed to film (SAGECREATION, MiNiChemi). Signal intensity for Dicer-2, H3K9me3, H3K27me3, H3K4me3 and Tublin was quantified using ImageJ.

For RNA quantification, total RNA was extracted, and genomic DNA was removed with DNase I (Roche, 04716728001). First-strand cDNA was generated with the SuperScript III Fist-Stand using random hexamers. The expression levels of RNAs were quantified on Rotor-Gene Q (QIAGEN) and normalized against GAPDH mRNA. PCR primers sequences were listed in Supplementary Table 3.

About 30 μg of total RNA isolated with Trizol was loaded into each lane of agarose gel and blotted onto membrane with Chemiluminescent Nucleic Acid Detection Module (Thermo Fisher) according to manufacturer’s instruction. RNA probes were labeled by *in vitro* transcription of plasmids with T7 RNA polymerase (Promega) in the presence of Biotin RNA labeling mix (Roche). The primers used were listed in Supplementary Table 3.

### Quantification and statistical analysis

Statistical parameters were reported either in individual figures or corresponding figure legends. Quantified data were in general presented as bar/line plots, with the error bar representing mean ± SEM, or boxplot showing the median (middle line), first and third quartiles (box boundaries), and furthest observation or 1.5 times of the interquartile (end of whisker). All statistical analyses were done in R. Wherever asterisks are used to indicate the statistical significances, *stands for p < 0.05; ** for p < 0.01, *** for p < 0.001 and NS for not significant.

## Data Access

All sequencing and processed files were deposited to Gene Expression Omnibus under accession number GSE134307. To review GEO accession GSE134307: Go to https://www.ncbi.nlm.nih.gov/geo/query/acc.cgi?acc=GSE134307, then enter token qzexqwwmbbujhol into the box. All software and the datasets used in this study are available from the corresponding author, Xiang-Dong Fu, upon reasonable request.

## ACKNOWLEDGEMENTS

We thank Dr. Yang Yu for providing S2 cells. This work was supported by grants from the National Key R&D Program of China (2016YFC0901702, 2016YFC09010002) and the National Natural Science Foundation of China (31871294, 31520103905) to R.S.C. and S.M.H., and NIH grants (HG004659, GM049369 and GM052872) to X.D.F.

## AUTHOR CONTRIBUTIONS

Conceptualization, X.D.F., R.S.C., and S.M.H.; Methodology development and data analysis, Y.J.H.; Generation of the GRID-seq data in S2 cells, X.L.; Experimental design and execution, Y.J.H., D.P.W. and S.H.W.; Data interpretation and discussion, Y.J.H., X.D.F., R.S.C., S.M.H., D.P.W, S.H.W, P.Z., J.Y.C., C.W.S., and D.H.L.; Paper writing, Y.J.H. and X.D.F.

